# Hierarchical Reconfiguration of Neurocognitive Task Set Representations Mediates Cognitive Flexibility

**DOI:** 10.64898/2026.01.20.700694

**Authors:** Stephanie C. Leach, Xitong Chen, Kai Hwang

## Abstract

Cognitive control organizes abstract contexts, stimuli, and actions into hierarchically structured representations. This organization supports flexible behavior but requires updating at multiple levels of the hierarchy, a process reflected in task switch costs. However, it remains unclear how updating differs across levels of abstraction and how these differences relate to behavior and neural representations. Here, we investigated the behavioral and neural sources of switch costs using fMRI and behavioral data from healthy adult participants (male and female). We employed a hierarchical control task that dissociates abstract context reconfiguration from more concrete task-set reconfiguration of stimulus-response mappings. We predicted that task sets, which incorporate sensory–motor mappings, would be more strongly influenced by feedforward inputs than higher-level contextual goals, which are more abstracted from immediate perceptual and motor demands. As predicted, subordinate rule switches were faster and were more strongly influenced by task-irrelevant perceptual changes, whereas context switches were slower and relatively insensitive to such interference. To characterize the neural basis of these effects, we quantified trial-to-trial reconfiguration of multivoxel activity patterns. Across the brain, larger pattern shifts predicted larger RT switch costs, linking representational reorganization to behavioral performance. Importantly, representational reconfiguration differed across hierarchical levels and anatomical systems. Subordinate rule updating was modulated by task-irrelevant perceptual features and expressed in distributed perceptual and motor networks, whereas context reconfiguration engaged the mid-lateral frontal cortex and was comparatively insulated from interference. Our results reveal how the hierarchical structure of neural representations supports flexible updating with interference-shielded contextual representations subserving behavioral control.

**Significance:** Daily activities often require simultaneously updating different tasks and thoughts. Driving requires maintaining a stable destination goal while rapidly updating motor plans (brake, accelerate, turn, etc.) in response to changing perceptual information (traffic lights, pedestrians, etc.). Although people perform such tasks with ease, it remains unclear how neural and cognitive representations are structured to respond to these flexibility demands. The present study suggests that sensory-motor plans prioritize flexibility by allowing greater influence from sensory inputs, which can create interference across brain networks when that input is task-irrelevant. Contextual information is insulated from this interference by representing contexts as distinctly as possible in the lateral prefrontal cortex, resulting in slower but more stable context switching.

## 1. Introduction

Everyday tasks require maintaining/updating multiple cognitive representations to support flexible, goal-directed behavior. For example, driving involves maintaining a destination goal while continuously updating route plans and switching between accelerating and braking in response to dynamic sensory inputs. These representations operate at different levels of abstraction. Task sets specify stimulus–response mappings and therefore likely incorporate perceptual and motor representations tied to sensory inputs, whereas contextual representations are more abstract and rely on internally maintained goals and memory (Badre & D’Esposito, 2007; Sakai, 2008). This organization introduces a core challenge: to support flexibility, control systems must update representations that differ in abstraction and sensitivity to feedforward sensory inputs. Understanding how this flexibility is achieved requires characterizing the representational structures that implement cognitive control (Badre et al., 2021; Monsell, 2003; Rogers & Monsell, 1995).

The structure of control representations encoding goals, contexts, task sets, stimuli, and actions shapes cognitive flexibility. Contextual representations, often associated with rostral mid-lateral prefrontal cortex (Badre & D’Esposito, 2007; Badre et al., 2009; Nee & Brown, 2013), are assumed to bias downstream processes to prioritize task-relevant information for hierarchical control (Miller & Cohen, 2001; Nee & D’Esposito, 2017). Such biasing can prioritize shielding representations from interference (Sakai, Rowe, & Passingham, 2002), while enabling flexible updating as contexts change and attention is redirected (Cellier, Petersen, & Hwang, 2022), including the selection of task-relevant rule–stimulus–response conjunctions (Kikumoto & Mayr, 2020). Flexibility may therefore depend on how abstract contexts organize representations that differ in their sensitivity to feedforward inputs (Schumacher & Hazeltine, 2016).

The structure of control representations can be probed by examining trial-to-trial reconfiguration (Bustos et al., 2024; Dykstra et al., 2022). Behavioral paradigms have traditionally examined reconfiguration via response time (RT) switch costs—the additional time it takes to make a response when a task changes from one trial to the next (Monsell, 2003; Rogers & Monsell, 1995; Wylie & Allport, 2000). Because internal representations constrain observable behavior, systematic variation in switch costs can reveal their underlying structure. We therefore hypothesize that sensitivity to feedforward interference depends on the extent to which control representations incorporate sensory–motor features. For task sets closely linked to sensory–motor features, task-irrelevant perceptual changes should modulate switch cost magnitude. In contrast, switch costs associated with shifting contexts specifying task sets should be less affected by task-irrelevant perceptual changes, reflecting greater abstraction from sensory–motor features.

Although switch costs provide a behavioral readout of reconfiguration processes, their neural basis remains unclear. Recent work suggests information is encoded in distributed activity patterns across neuron populations, voxels, or electrodes (Ebitz & Hayden, 2021; Mur, Bandettini, & Kriegeskorte, 2009). Under this population view, cognitive flexibility entails transforming one activity pattern into another. The magnitude of this transformation can be quantified as the multivariate similarity between activity patterns on successive trials, where larger dissimilarity reflects greater representational reorganization and predicts greater RT switch costs (Qiao et al., 2017). Building on these findings, we hypothesize that task sets incorporating sensory–motor features are encoded in more similar patterns and therefore undergo smaller reconfiguration during switching. In contrast, contextual representations, being more abstract, are encoded in more distinct patterns to reduce interference, and consequently require greater reconfiguration during switching.

Given these hypotheses, our study pursued two goals. First, we leveraged RT switch costs to determine how hierarchical task structure and feedforward perceptual inputs influence the reconfiguration of distinct hierarchical representations. Second, we quantified trial-to-trial shifts in multivoxel activity patterns (henceforth “multivoxel pattern shifts”) to examine neural correlates of representational reconfiguration and perceptual influence. Results confirmed that larger pattern shifts predicted larger RT switch costs. Moreover, representational reconfiguration differed across hierarchical levels in its sensitivity to perceptual input. Increased feedforward perceptual influence modulated subordinate rule selection but not context selection. These findings suggest that contextual representations organize task sets and rule–stimulus–response conjunctions in a representational format that is relatively insulated from task-irrelevant perceptual information.

## 2. Methods

### 2.1. Participants

Participants were recruited for two experiments, experiment 1 involved MRI scanning, and experiment 2 was a behavioral only experiment conducted to replicate in-scanner behavioral effects. For experiment 1, we recruited 73 healthy participants (53 female, 20 male, *M*_age_ = 22.31, *SD*_age_ = 4.45, *Range*_age_ = 18-35 years). Out of these 73 participants, 10 were excluded for poor behavioral performance (accuracy < 75% for at least one condition), and 9 were excluded for excessive movement artifacts (mean framewise displacement > 0.4 mm across runs). After exclusion, we included data from 54 participants (39 female, 15 male, *M*_age_ = 22.38, *SD*_age_ = 4.87, *Range*_age_ = 18-35 years). These data were also published in our previous study (Chen et al., 2024). For experiment 2, we recruited 65 healthy participants (55 female, 9 male, 1 non-binary, *M*_age_ = 18.63, *SD*_age_ = 0.71, *Range*_age_ = 18-21 years). Out of these 65 participants, 8 were excluded for poor behavioral performance (accuracy < 75% for at least one condition). After exclusion, we included data from 57 participants (50 female, 7 male, *M*_age_ = 18.63, *SD*_age_ = 0.72, *Range*_age_ = 18-21 years). Both experiments were approved by the University of Iowa Institutional Review Board. All participants were recruited from the University of Iowa and surrounding area to participate. All participants had normal or corrected to normal visual acuity and color vision, were right-handed, and with no history of epilepsy, psychiatric, or neurological conditions. All participants gave written informed consent. The experiments were conducted in compliance with the ethical principles expressed in the Declaration of Helsinki.

### 2.2. MRI data Acquisition

Imaging data were collected at the Magnetic Resonance Research Facility at the University of Iowa using a 3T GE SIGNA Premier scanner with a 48-channel head coil. Structural images were acquired using a multi-echo MPRAGE sequence (TR = 2348.52ms; TE = 2.968ms; flip angle = 8°; field of view = 256*256; 200 sagittal slices; voxel size = 1mm^3^). Functional images were acquired using an echo-planar sequence sensitive to blood oxygenated level-dependent (BOLD) contrast (Multiband acceleration factor = 2; no in-plane acceleration; TR = 1800ms; TE = 30ms; flip angle = 75°; voxel size = 1.718 mm*1.718mm *2.5 mm).

### 2.3. MRI Data Preprocessing

All fMRI data were preprocessed using fMRIPrep version 20.1.1 (Esteban et al., 2019) to reduce noise and transform data from subject native space to the ICBM 152 Nonlinear Asymmetrical template version 2009c for group analysis (Fonov et al., 2011). Preprocessing steps include bias field correction, skull-stripping, co-registration between functional and structural images, tissue segmentation, motion correction, and spatial normalization to the standard space. We did not perform any spatial smoothing and did not apply conventional nuisance regression. Instead, noise suppression was implemented during within the GLMsingle procedure (Prince et al., 2022). GLMsingle combines voxel-wise GLM estimation with automated denoising procedures from GLMdenoise (Kay et al., 2013), which uses cross-validation to identify and apply the optimal combination of nuisance regressors. Nuisance regressors, motion estimates, and principal components were learned to best improve cross-validation performance from whole-brain regions of interest. The framework also estimates single-trial beta weights while correcting for temporal autocorrelation and optimizing the hemodynamic response function on a voxel-wise basis (see section 2.7). This approach provides improved sensitivity and reliability for our trial-by-trial transition analyses.

### 2.4. Experimental Design

In experiment 1, participants underwent an fMRI scanning session while performing a hierarchical cognitive control task. Before entering the scanner, all participants completed a tutorial and a practice session. Once participants achieved at least 80% on the practice, the experiment’s scanning session began. We collected eight runs of functional scans, with each run consisting of 51 task trials. This resulted in a total of 408 task trials across all eight runs. Each functional run lasted approximately 6.5 minutes. In experiment 2, participants performed the same hierarchical cognitive control task behaviorally with the same training procedure, trial timing, block structure, and number of trials. The purpose was to replicate the behavioral effects we observed in scanner.

### 2.5. Behavioral Tasks

To study the representation structure for flexible hierarchical cognitive control, we developed a hierarchical cognitive control task (Chen et al., 2024). The task implements a three-level hierarchy in which texture defines context, context determines the relevant feature dimension (color or shape), and the feature dimension maps onto a response rule.

#### 2.5.1. Task structure and trial sequence

In this task, each trial began with a cue object presented for 500 milliseconds, followed by an image of either a face or a scene presented for 2000 milliseconds (**Figure 1A**). The inter-trial interval (ITI) was jittered between 1.5 to 10.5 seconds with an average duration of 4.25 seconds. Depending on the cue object presented before the image, participants made a yes/no judgment on one of two response rules: (1) was the image a face? (2) was the image a scene? The luminance of probe images (faces/scenes) was equalized using the SHINE toolbox (Willenbockel et al., 2010). Face images included a scrambled background to control for low level spatial frequency to match the scene images. Probe images were randomly selected from a set of 50 unique faces and 50 unique scene images. Finally, we ensured that adjacent trials never used the exact same face or scene image.

**Figure 1.**
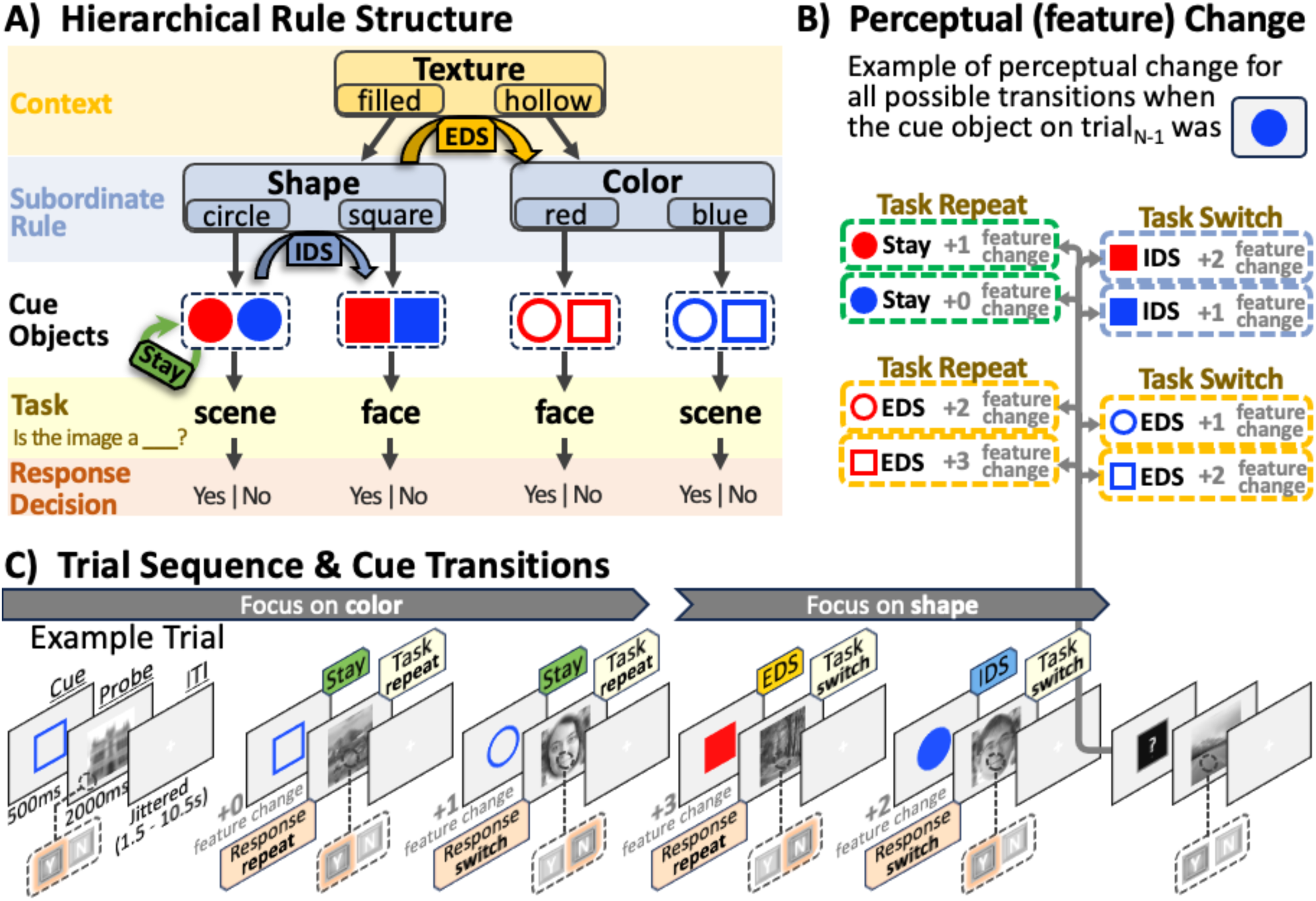
Hierarchical control task design. **A)** Depiction of the task rule hierarchy embedded within the 8 potential cue objects. Context switch (EDS) trials reflect switching between the shape and color rule, or switching between a filled and hollow cue object. Subordinate rule switch (IDS) trials reflect repeating context, but switching between either circle and square or red and blue, depending on the context rule (shape or color). Finally, Stay trials are trials where both context and subordinate rule repeat. **B)** More detailed depiction of perceptual change showing eight possible transitions following a trial with a filled, blue, circle cue object. Across trials, 0-3 features could change. **C)** Example trial sequence: a cue object is presented for 0.5 seconds, followed by an image of either a face or a scene for 2 seconds, and a jittered ITI (range: 1.5-10.5 seconds; average: 4.25 seconds). Example cue transitions across trials depicts how perceptual change is operationalized. The first Stay trial shows a cue transition with 0 features changing. The second Stay trial shows a cue transition where one cue feature changes (shape, a task-irrelevant feature in this example). The EDS trial shows a cue transition where all three cue features change. The IDS trial shows a cue transition where two features change (shape, task-relevant; color, task-irrelevant). Example face images are authors on this manuscript.

#### 2.5.2. Hierarchical rule structure

To determine which response rule to use on a given trial, participants had to integrate three sources of information associated with three perceptual features of the cue object: texture, shape, and color (**Figure 1B**). The cue’s texture determined whether the response rule was signaled by shape or color. If the texture of the cue object was filled in, the shape of the object (square or circle) would indicate which rule they needed to use (square being the face rule, “is the image a face?”, and circle being the scene rule, “is the image a scene?”). Alternatively, if the texture of the cue object was hollow (i.e., only an outline), the color of the object (red or blue) would indicate which rule they needed to use (blue being the face rule and red being the scene rule). Half of the participants completed a counterbalanced version of this task with alternative yes/no response key mappings, and in which the top level of the hierarchy was flipped (i.e., hollow texture cues shape and filled texture cues color).

#### 2.5.3. Hierarchical rule switching

This task design yields three types of trial-to-trial transitions that differ in the level of hierarchical reconfiguration required: Stay, intra-dimension switch (IDS), and extra-dimension switch (EDS). On Stay trials, participants repeated the same rule as the previous trial, preserving both the texture-defined context and feature-to-rule mapping (e.g., which color or shape indicates the face task). On IDS trials, participants switched between task-relevant features (shape vs. color) within the same texture context. On EDS trials, they updated the highest-level context by switching texture cues (filled vs. hollow), thereby changing which feature signaled the response rule.

#### 2.5.4. Trial-to-trial transitions, perceptual, and motor factors

In addition to hierarchical rule switching, the task design allowed us to quantify trial-to-trial perceptual and motor influences on performance. Specifically, (1) whether the texture, shape, or color of the cue object repeated from the previous trial, (2) how many cue object features changed (range: 0-3; **Figure 1C**) and (3) how these factors interacted with rule switching, (4) whether the motor response repeated, (5) whether the targeted, relevant feature repeated or switched from the previous trial. Here, previous target feature refers to the specific cue feature (i.e., red, blue, circle, or square) that defines the task rule. For example, if the current subordinate rule is the color rule and the current cue object is red, then the targeted, relevant feature is “red”. If on the next trial the current subordinate rule is the shape rule, but the cue object is still red, then the targeted relevant feature from the previous trial repeated even though the subordinate rule switched. As described in detail below, we used linear regression to quantify the influence of each factor on response times and reconfiguration on voxel activity patterns.

#### 2.5.5 Behavioral (Response Time) Switch Costs

To quantify behavioral costs associated with hierarchical task reconfiguration, we analyzed trial-wise response times using linear mixed-effects regression. This approach allowed us to estimate the contributions of hierarchical rule switching, perceptual feature transitions, and motor processes to performance costs while accounting for repeated measurements within subjects.

### 2.6. Trial-to-Trial Shifts in Multivoxel Activity Patterns

The goal of this analysis was to characterize how neural activity patterns that instantiate cognitive task representations are reconfigured from trial to trial. To this end, we quantified trial-to-trial changes in multivoxel BOLD activity patterns and examined how these representations shift from trial-to-trial (hereafter described as “multivoxel pattern shifts” or “pattern shifts”; Fig. 2A). The key underlying assumption here, in line with the population representation framework (Mur et al., 2009; Kriegeskorte et al., 2008), is that the degree of multivoxel pattern shifts can be used as an index of the degree of representational shifts. Briefly, larger representational shifts should result in larger multivoxel pattern shifts while smaller representational shifts should result in smaller multivoxel pattern shift; however, interference arising from small representational distances can also cause larger multivoxel patterns shifts under certain conditions eliciting this greater interreference, such as task-irrelevant changes (Fig. 2B). Using the same analytical logic as our behavioral switch cost analysis, we fitted regression models to pattern shifts to determine how reconfiguration was influenced by hierarchical task structure as well as perceptual and motor factors. This analysis included a multistep pipeline consisting of trial-wise activity estimation, computation of multivariate pattern similarity, and regression-based modeling that paralleled to the behavioral switch-cost analysis.

**Figure 2.**
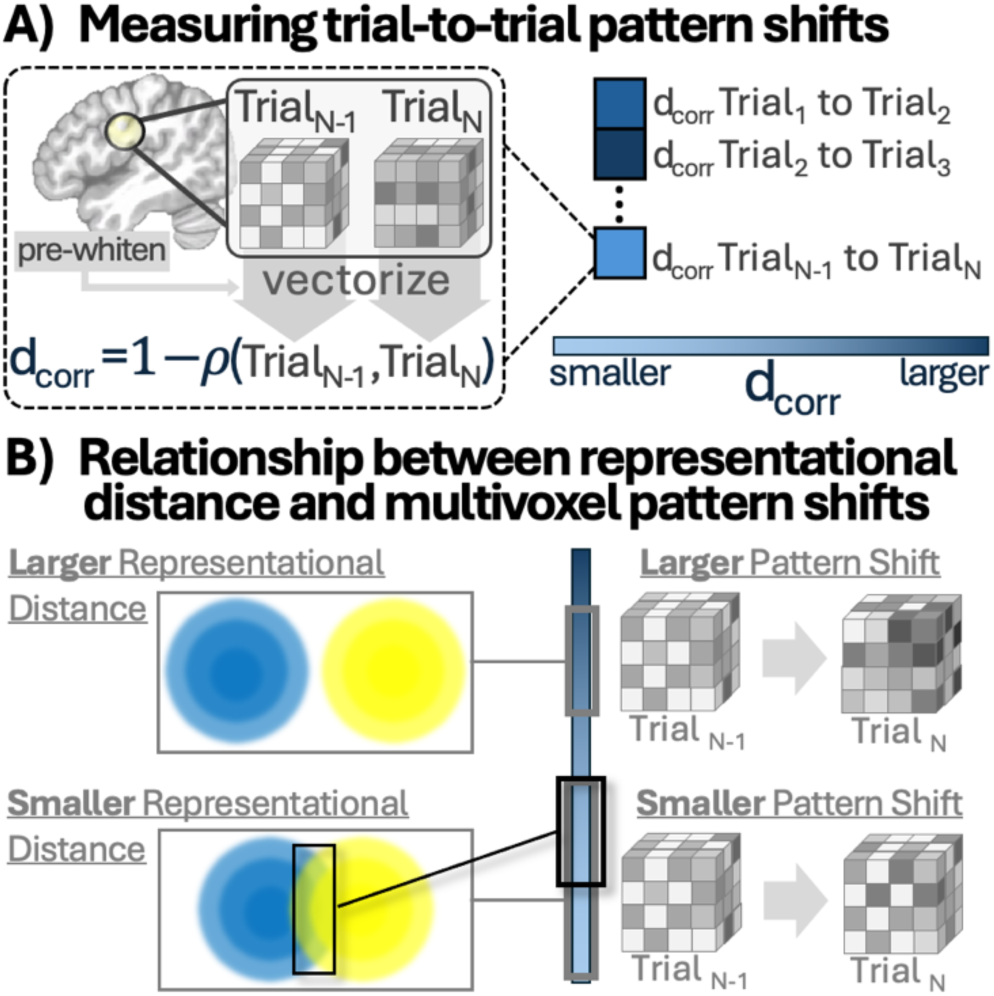
Multivoxel Pattern Shifts Method. **A)** Depiction of how pattern shifts are calculated on adjacent trial multivoxel activity patterns. A greater change in multivoxel activity patterns is associated with a greater correlation, or similarity. **B)** Depiction of relationship between pattern shifts and representational distance. In general, a larger representational distance should result in a greater pattern shift while a smaller representational distance should result in a smaller pattern shift; however, interference between representations can also cause greater pattern shifts.

#### 2.6.1. Trial-wise activity estimate

Preprocessed fMRI time-series (outputs of fMRIPrep) were first submitted to GLMsingle (Prince et al., 2022) to estimates trial-wise beta estimates. GLMsingle fits a general linear model (GLM) with one regressor per trial and uses cross-validation to improve estimation. Specifically, voxel-wise noise regressors are derived from the data and evaluated on held-out runs, and a regularized (ridge) regression model is fit with the regularization parameter (λ) selected via cross-validation. This approach reduces collinearity among adjacent trial estimates and improves the stability of single trial β values. In addition, GLMsingle estimates a voxel-specific hemodynamic response function (HRF), allowing the model to account for regional differences in neurovascular coupling rather than assuming a fixed HRF across the brain. The model was implemented with a 0.5 s event duration corresponding to cue onset, because there was no temporal jitter between the cue and probe stimuli, the estimated responses reflect activity across the entire trial. A library of gamma functions with varying shapes was used to identify the best-fitting HRF for each voxel to fully capture responses to cue and probe objects. At the end, this procedure yielded one amplitude estimate per trial per voxel. To ensure that variability in RT did not drive the multivoxel pattern shifts, we regressed each trial’s RT from its corresponding amplitude estimate on a voxel-by-voxel basis before further analyses.

#### 2.6.2. Multivoxel Pattern Shifts

Using these RT-corrected trial-wise maps, we then calculated multivariate distances between trial-wise activity patterns. This was done with an 8 mm radius volumetric searchlight analysis, restricting to gray matter voxels. Within each searchlight sphere, signals were first “whitened” by applying the noise covariance matrix (equation 1) estimated across all trials and voxels in that sphere to remove shared noise correlations (equation 2) before measuring pattern shifts (Walther et al., 2016).

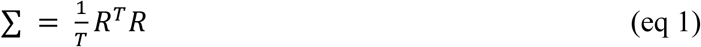

Where R is the model residuals matrix of size T x P. Note, T is the number of time points and P is the number of voxels. The covariance matrix, Σ, is size P x P.

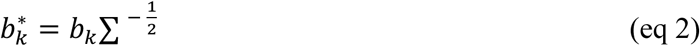

Where b_k_ is a searchlight sphere multivoxel activity pattern of size P corresponding to one condition, k, of the possible K conditions.

This procedure removes voxel-wise correlations attributable to shared noise. We then computed the correlation distance (equations 3 and 4) between consecutive trials:

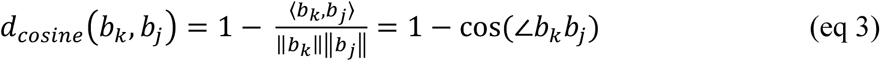

Where b_j_ is a searchlight sphere multivoxel activity pattern of size P corresponding to one condition, j, of the possible K conditions.

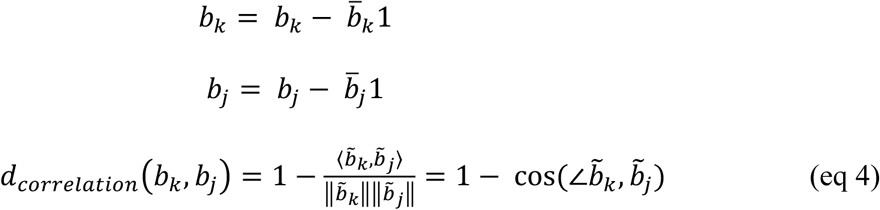

Where 1 is a row vector of ones of size 1xP and *b̅* is the voxel mean.

Correlation distance was chosen over Euclidean or Mahalanobis metrics because we were primarily interested in changes in the neural representation geometry rather than overall amplitude shifts. This procedure yielded, for each voxel, a time series of pattern shifts values reflecting trial-to-trial representational reconfiguration.

### 2.7. Statistical Analyses

#### 2.7.1 Behavioral Model overview

The primary goal of the behavioral linear mixed effects model (Bates et al., 2015) was to identify factors that influence response time switch costs during hierarchical cognitive control. Specifically, the model tested how performance costs varied as a function of (1) the level of hierarchical rule reconfiguration (Stay, IDS, EDS), (2) trial-to-trial changes in perceptual cue features, and (3) interactions between hierarchical switching and perceptual change. Additional regressors were included to control for known effects of motor repetition, task repetition, and error-related adjustments.

#### 2.7.2. Behavioral preprocessing and dependent variable (z-scored response times)

The dependent variable was trial-wise response time, normalized as z-scored response times (zRTs). Specifically, we first normalized each participants’ RT within each experimental block and response modality (i.e., yes versus no response). When normalizing RTs we did not include error or post-error trials when calculating the mean or standard deviation because of extensive work showing error and post-error slowing effects (McClure, Berns, & Montague, 2003; Rabbitt, 1966). We then applied the following exclusion criteria: (1) the first trial of each run, (2) any zRTs exceeding 3 standard deviations from the mean, (3) any trials with missing responses, (4) any post-error trials that were also erroneous.

#### 2.7.3. Primary behavioral model predictors

For the model regressors, we included the following fixed effects. First, we included hierarchical task switching (Stay, IDS, EDS) as a categorical predictor to capture the effects of abstract rule structures on performance cost. To determine how perceptual representations of the cue attributes influence switch costs beyond abstract rules, we incorporated several perceptual factors: cue texture, color, and shape, along with the number of features that changed from the previous trial (range: 0-3).

We further included an additional binary regressor indicating whether the target feature repeated or switched from the previous trial. For example, if the relevant feature on trial N-1 was color (red), the target feature was coded as “repeated” if the cue on trial N was also red, and “switched” if the cue was a different color (e.g., blue). This regressor indexed how the previous trial’s task-relevant perceptual representation influences current trial performance.

Critically, we included an interaction between hierarchical task switching and the number of perceptual features that changed from the previous trial. This critical interaction tested our hypotheses about flexibility prioritization. If lower-level rule representations prioritize flexibility, we expected a significant positive interaction between IDS trials and feature change count, given that a greater number of perceptual features changing meant task-irrelevant features were also changing. Conversely, if contextual representations prioritize stability, we expected no significant interaction between EDS trials and feature change.

#### 2.7.4. Behavioral model control variables and conjunction related interactions

To account for known influences on response times, we included several control variables: task-rule repetition vs. switch, yes/no response repetition vs. switch, and probe-image repetition vs. switch. We also controlled for trial-by-trial performance fluctuations by including separate regressors for post-error trials and error trials to account for post-error slowing effects.

Beyond controlling for nuisance variance, the model incorporated a set of theoretically motivated interaction terms to replicate established task-switching effects. In particular, prior work suggests that flexible behavior is supported by conjunctive representations that integrate task rules, stimuli, and responses, and that the strength of such conjunctions can be indexed indirectly through partial switch costs (Grzyb & Hubner, 2013; Kikumoto & Mayr, 2020; Kleinsorge & Heuer, 1999). Partial switch costs arise when some, but not all, components of a task representation change across trials, requiring unbinding and rebinding of the prior conjunction (Li et al., 2019). Therefore, we further included a theoretically motivated interaction term: task-rule repetition with response repetition. This interaction terms tests the canonical partial switch costs, characterized by faster responses on full repetition and full switch trials relative to partial repetition/switch trials (Grzyb & Hubner, 2013; Kleinsorge & Heuer, 1999).

Because this interaction term indexing conjunction related effects could differ as a function of hierarchical context, we ran a second mixed effects regression model where we replaced hierarchical switch type (Stay/IDS/EDS) with a binary regressor indicated in context repeated or switched. This further allowed us to include two-way interactions between (1) context repeat and task repeat and (2) context repeat and response repeat, as well as the 3-way interaction of primary interest: context repeat by task repeat by response repeat. If contextual representations organize lower-level task sets, i.e., task rule–stimulus–response associations, then partial switch costs should primarily occur within contexts and be weaker across contexts. In other words, we should observe a significant three-way interaction between context, task, and response that is driven by the existence of (1) a significant two-way interaction between task and response within context, but (2) a non-significant two-way interaction between task and response when switching across context.

#### 2.7.5. Behavioral model random effects structure

We treated subject as a random intercept and estimated random slopes for each individual cue feature (color, shape, and texture) to capture individual differences in sensitivity to these perceptual factors. Due to the complexity of the model, we kept the random effects structure as simple as possible to avoid convergence issues. The full model was specified as:

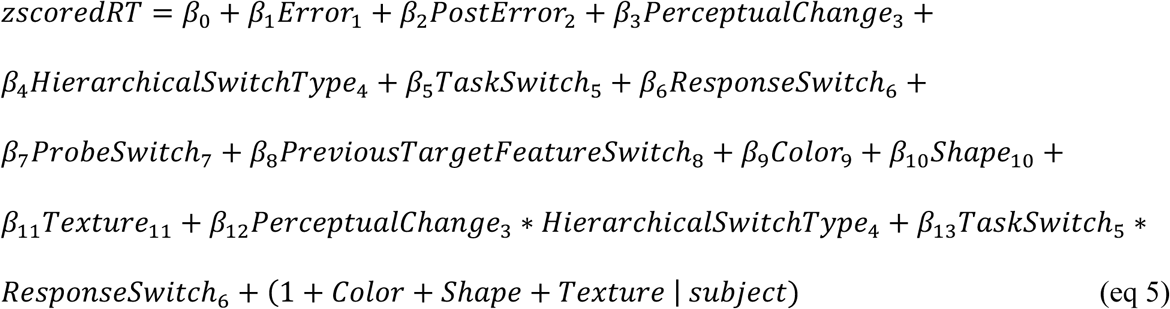

#### 2.7.6. Behavioral (response time) supplementary analysis

It is important to note a subtle difference between IDS and EDS trials regarding trials where two features changed. For IDS, a two-feature change always reflects a change in both a task-relevant and task-irrelevant feature. In contrast, a two-feature change on EDS trials does not always mean a change in one relevant and one irrelevant feature. Sometimes a two-feature change on EDS trials means two relevant features change and no task-irrelevant features change. Therefore, we ran an additional follow-up paired samples t-test where we ensure EDS trials where two features changed were restricted to only trials where one feature was task-relevant and the other was task-irrelevant. We then contrasted these trials with trials where only one task-relevant feature changed.

#### 2.7.7 Regression model for pattern shifts

For each subject, the pattern shift time series was entered as the dependent variable in a linear regression model closely paralleling the behavioral model described in above. The primary predictors included hierarchical task-switching condition (Stay, IDS, EDS), perceptual feature-change variables, motor repetition, error and post-error indicators, and their theoretically motivated interactions. Unlike the behavioral model, regressors indexing the absolute texture, color, or shape of the current trial’s cue were not included because the dependent variable was a measure of dissimilarity between the current trial and previous trial, whereas the behavioral model’s dependent variable was a measure of the magnitude of the current trial’s performance cost.

To further control for potential RT effects on the BOLD response (Mumford et al., 2024), which was used to calculate pattern shifts, we added a regressor reflecting the magnitude of trial-to-trial changes in zRT (difference in zRT across adjacent trials) and a regressor reflecting the magnitude of the current trial’s zRT. The model was specified as:

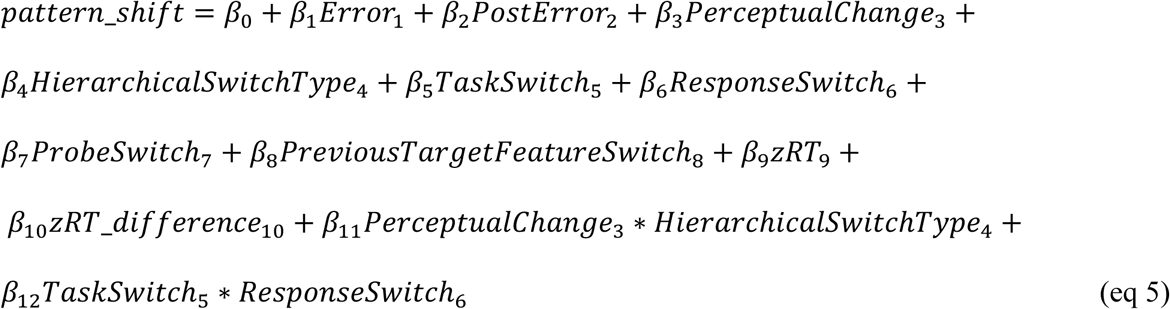

#### 2.7.8 Group analysis of pattern shifts

This procedure yielded, for each subject, a whole-brain beta map of each regressor that was then entered into a one-sample t-test across participants. To control the family-wise error rate for multiple comparisons, we employed AFNI’s 3dClustSim using the residuals from the GLM procedure (Cox et al., 2017). First, spatial autocorrelation of the residuals was estimated using the autocorrelation function (ACF), producing three ACF parameters that characterize the spatial smoothness of the data. These parameters were then supplied to 3dClustSim to estimate the minimum cluster size for a corrected cluster-level threshold. Significant clusters for parametric modulation are based on these calculated minimum cluster sizes of 22 voxels (cluster p-value threshold of .001 and a voxel-level p-value threshold of .005).

### 2.8. Behavioral Switch Cost and Multivoxel Pattern Shift Predictions

Our framework posits that hierarchically organized representations differ in their level of abstraction and their sensitivity to sensory input. Task sets, which implement task rule-stimulus–response mappings, are more tightly coupled to immediate sensory-motor plans, whereas contextual representations are more abstracted from these immediate sensory-motor plans. These differences are expected to influence both behavioral switch costs and trial-to-trial changes in multivoxel activity patterns (pattern shifts).

Our predictions follow the population representation framework in which multivoxel pattern similarity is taken to reflect representational similarity (Kriegeskorte, Mur, & Bandettini, 2008; Mur et al., 2009). Under this framework, different task representations evoke distinct distributed activity patterns, and greater dissimilarity between patterns indexes less similar underlying representations (Cox & Savoy, 2003; Kriegeskorte et al., 2008). Prior work has shown that both categorical and parametric differences in representational content, including abstract task variables, are reflected in systematic variation in multivoxel patterns (Garvert et al., 2017; Park et al., 2020). Accordingly, we used multivoxel pattern shifts as an index of changes the underlying representations engaged by the task.

For representations that incorporate sensory-motor plans, we predict increased sensitivity to interference from task-irrelevant perceptual changes. Specifically, we expect the reconfiguration of these representations to be more susceptible to interference from task-irrelevant information due to (1) smaller representational distance and (2) greater sensory-driven influence driving representational reconfiguration. We test this prediction by examining the interaction between lower-level rule reconfiguration (IDS) and the number of stimulus features (texture, color, shape) that change across trials. On IDS trials, the contextual texture repeats while the relevant feature (color or shape) switches. When only one feature changes, only the task-relevant dimension is updated; however, when multiple features change, both relevant and irrelevant dimensions are updated. We predict that changes in task-irrelevant features will increase both switch costs and pattern shifts, reflecting greater feedforward influence.

For EDS trials, which require switching the context, we predict the opposite pattern. Contextual representations are abstracted away from immediate sensory-motor plans, and therefore we hypothesize that they are separated by larger representational distances and less influenced by trial-to-trial changes in perceptual and motor factors. Accordingly, the interaction between feature change and switch type should be weaker for EDS than for IDS or Stay trials. In addition, because context switching involves transitions between more distinct task sets, we expect larger overall pattern shifts for EDS relative to IDS.

### 2.9. Code and Data Availability

All raw data are publicly accessible on OpenNeuro (https://openneuro.org/datasets/ds005600/). Any preprocessing/analysis scripts used in this study can be found on GitHub (https://github.com/HwangLabNeuroCogDynamics/Neurocognitive_TaskSets).

## 3. Results

We analyzed both behavioral (z-scored RTs) and neural reconfiguration (pattern shifts) measures using a common regression framework to identify factors influencing switch costs and multivoxel pattern reconfiguration during hierarchical control. Specifically, we examined how hierarchical level (context vs. subordinate rule switching) and perceptual factors modulated behavioral performance and multivoxel pattern shifts. In addition, we tested interactions between context and rule–stimulus–response conjunctions (Kikumoto and Mayr, 2020). More specifically, we leverage partial switch costs as an indirect measure of conjunction strength, to test whether different contexts are split into distinct representational subspaces, within which context-dependent conjunctions are embedded.

### 3.1. Distinct hierarchical level representations interact differently with task-relevant and task-irrelevant perceptual influences during reconfiguration

Traditionally, a hierarchical representation structure has been assessed by comparing the magnitude of switch costs across different levels of hierarchical rule switching (Fig. 3A, top panel). Larger switch costs when context switches relative to subordinate-rule switches have been taken as evidence of an underlying hierarchical representation structure (Cellier et al., 2022; Chen et al., 2024). The present dataset displays these expected switch cost effects of larger z-scored RT switch costs for context switches (EDS) as compared to both subordinate rule switches (IDS; *t*(53)=8.010, *p*<.001), and rule repeats (*t*(53)=18.128, *p*<.001), as well as subordinate rule switches (IDS) as compared to rule repeats (Stay; *t*(53)=11.195, *p*<.001; Fig. 3B). Switch costs that increase with reconfiguration of increasingly superordinate rules suggest an underlying hierarchical organization of task representations in the present dataset.

**Figure 3.**
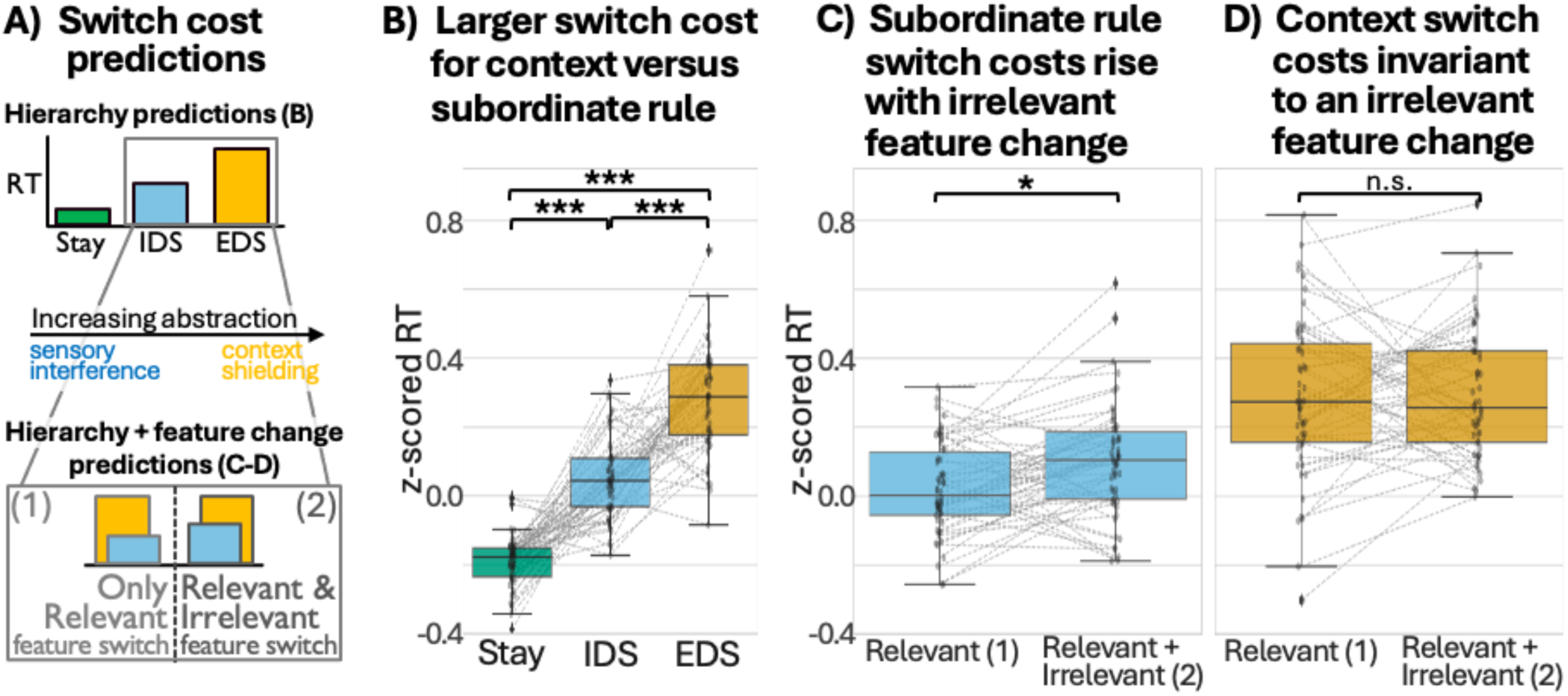
Response Time Switch Costs. **A)** Schematic of predicted switch cost patterns for a hierarchical representation structure (top) and a more fine-grained analysis incorporating influences from task-relevant and task-irrelevant features (bottom). **B)** Switch cost comparisons across hierarchical rule conditions (Stay, IDS, EDS), showing increasing switch cost magnitude with increasing abstraction. Significance is based on two-tailed paired sample t-tests with z-scored RTs. **C-D**) Switch cost effects from the interaction between hierarchical rule (Stay, IDS, EDS) and the number of features that changed from the previous trial (perceptual change). Switch costs increased with task-irrelevant feature changes during subordinate rule switching (**C**), but not during context switches (**D**). Significance is based on post-hoc comparisons between estimated marginal means for hierarchical rule (Stay/IDS/EDS) and perceptual change. *** means p < .001, ** means p < .01, * means p < .05, n.s. means p >.05.

However, focusing solely on the magnitude of switch costs across hierarchical levels neglects critical interactions between hierarchical task structure and the processing of task-relevant versus task-irrelevant perceptual information across trials. In particular, trial-to-trial changes in cue features can influence performance depending on whether task-relevant or task-irrelevant cue features repeat or switch. Consider two examples. First, a cue may change from a red circle to a red square, in which only the task-relevant feature (shape) changes while the task-irrelevant feature (color) repeats. Second, a cue may change from a red circle to a blue square, in which both task-relevant and task-irrelevant features change. At present, it remains an open question as to whether these two example transitions differentially interact with subordinate rule and contextual reconfiguration processes.

Arguably, sensitivity to task-irrelevant perceptual change should provide a behavioral marker of how representations are organized. If representations are closely tied to sensory–motor features, changes in task-irrelevant dimensions should increase interference and slow performance. In contrast, more abstract representations should be less influenced by such changes. In other words, a benefit of a hierarchical representation structure is that contextual representations can be abstracted away from immediate sensory-motor plans, insulating superordinate levels from feedforward interference at the cost of overall greater performance costs associated with contextual reconfiguration. To test this, we conducted more fine-grained analyses examining how trial-to-trial perceptual feature changes interact with hierarchical rule reconfiguration (Fig. 3A, bottom). Switch trials (IDS and EDS) were subdivided based on whether only the task-relevant feature changed (one feature changed) or both task-relevant and task-irrelevant features changed (two feature changed).

Consistent with our predictions (Fig. 3A-bottom), if subordinate rules prioritize flexibility by allowing greater feedforward influence from perceptual input, then subordinate rule reconfiguration should be more susceptible to interference from changes in task-irrelevant cue features. Supporting this prediction, trial-level mixed-effects models (with random intercepts for subjects) revealed positive slopes for the number of features changing across trials (perceptual change) for both rule repeat (Stay; *β* = 0.151, *SE* = 0.017, *CI* = [0.117, 0.184]), and subordinate rule switch (IDS; *β* = 0.072, *SE* = 0.023, *CI* = [0.026, 0.118]) and a negative slope for context switch (EDS; *β* = -0.051, *SE* = 0.023, *CI* = [-0.095, -0.007]).

Follow-up contrasts confirmed that these positive slopes were associated with significantly slower response times when more cue features changed for both rule repeats (Stay; zero versus one; *mean difference* = 0.151, *t*(19588) = 8.835, *p* < .001; degrees of freedom reflect trial-level mixed-effects analyses, Satterthwaite’s approximation (Kuznetsova, Brockhoff, & Christensen, 2017), with all appropriate trials included, and subjects modeled as random intercepts), and subordinate rule switches (IDS; one versus two; *mean difference* = 0.072, *t*(20227) = 3.063, *p* = .013; Bonferroni corrected (Benjamini & Hochberg, 1995); Fig. 3C). This indicated that, as predicted, the additional task irrelevant feature change incurred an additional switch cost, i.e., slower response times, for subordinate rule switches. Moreover, for EDS trials there was no significant increase in response times when more cue features changed for context switch (EDS) trials (one versus two, *mean difference* = 0.051, *t(20888)* = -2.266, *p* = .141; Fig. 3D; one versus three, *mean difference* = 0.102, *t(20888)* = -2.266, *p* = .141, two versus three, *mean difference* = 0.051, *t(20888)* = -2.266, *p* = .141). This suggests that context reconfiguration is better shielded from potential feedforward perceptual interference from task-irrelevant feature changes. Note, because a two-feature change on EDS trials did not always mean both a task-relevant and task-irrelevant feature change, we ran a supplementary post-hoc paired sample t-test where we ensured this was the case (see methods section *Behavioral (response time) supplementary analysis* for more details). Critically, this analysis also found no difference in the magnitude of zRT switch costs between one versus two features changing (paired-sample t-test *t*(53) = -0.010, *p* = .797).

Finally, we found a significant interaction between these perceptual change slopes for the different rule reconfiguration conditions (i.e., IDS versus EDS), indicating that this slope is significantly more positive for IDS trials as compared to EDS trials (*β* = 0.123, *t(20860)* = 4.123, *p* < .001). To ensure the effects reported above were not driven by other trial-to-trial factors, we included several control regressors in the model, including task switches (*β* = 0.018, *t(*20880) = 0.506, *p* = .613), switching responses (*β* = 0.019, *t(*20880) = 0.683, *p* = .495), switching probe images (*β* = 0.027, *t(*20880) = 1.071, *p* = .284), error trials (*t(20,870)* = 10.504, *p* < .001) and post-error trials (*t(20,820)* = 2.359, *p* = .018), as well as switching the task-defining (target) feature from the previous trial (*t(20,820)* = 2.359, *p* = .018).

All in all, these results support our hypotheses on cognitive flexibility. Specifically, the degree of perceptual change in the cue object interacts with rule reconfiguration differently depending on which hierarchical level is being reconfigured. Subordinate rule representations support flexibility, i.e., faster switching, at the cost of greater perceptual interference during reconfiguration. Conversely, contextual representations are abstracted away and therefore less influenced by perceptual interference during reconfiguration, but show greater overall performance costs, i.e., slower switching, as a result.

### 3.2. Hierarchical reconfiguration and feedforward perceptual inputs jointly shape multivoxel pattern shifts

To investigate potential neural underpinnings of these z-scored RT switch cost effects, we conducted a parallel analysis using trial-to-trial changes in neural representational patterns (“multivoxel pattern shifts”) as a neural analogue of switch costs. We applied the same regression model used for the behavioral switch cost analysis to predict multivoxel pattern shifts. We included trial-wise z-scored RT and its derivative as additional predictors to control for potential effects of RTs on neural dissimilarity patterns (Mumford et al., 2024).

Based on prior work implicating the middle frontal gyrus (MFG) in contextual encoding and hierarchical cognitive control (Badre & D’Esposito, 2007; Badre & Nee, 2018; Nee & D’Esposito, 2016), we predicted that context reconfiguration (EDS) would elicit a larger pattern shift in MFG than subordinate rule reconfiguration (IDS). In parallel with behavioral findings, we further predicted that EDS reconfiguration would be relatively insensitive to perceptual interference. In contrast, we predicted that IDS reconfiguration would interact with perceptual interference (Fig. 4A). Specifically, we predicted that pattern shifts in MFG during EDS trials would be unaffected by the number of perceptual cue features changing across trials. In contrast, we predicted that premotor and visual attention regions would show larger pattern shifts on IDS trials when a greater number of perceptual features changed. Presumably, the latter would reflect both the encoding of subordinate rules in premotor regions (Badre & Frank, 2012; Nee & Brown, 2013) and increased processing of task-irrelevant perceptual information during subordinate rule reconfiguration.

**Figure 4.**
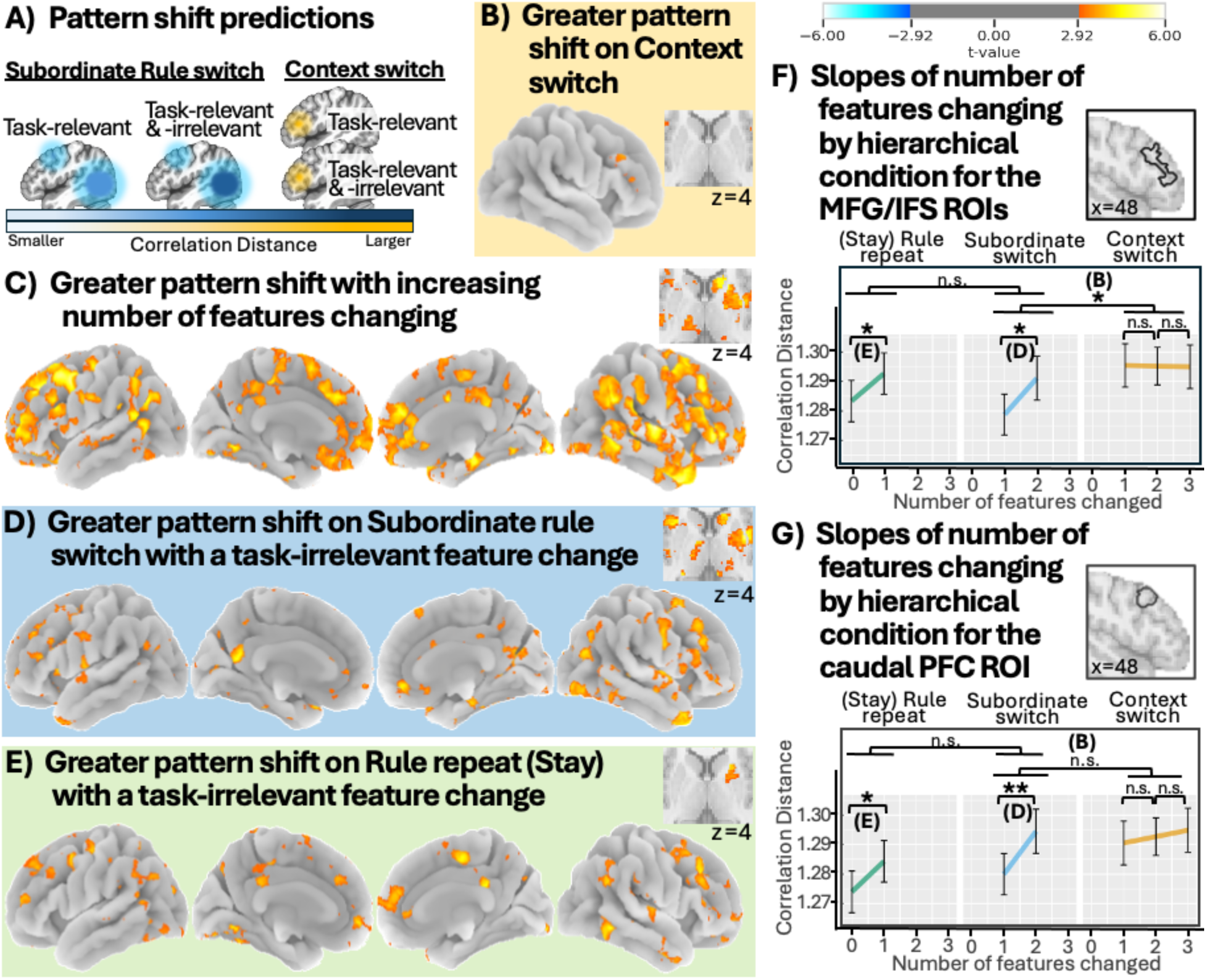
Reconfiguration of Multivoxel Patterns Across Hierarchical Levels Shows Differential Sensitivity to Perceptual Change. **A)** Schematic of expected effects for context versus subordinate rule reconfiguration. **B)** Observed main effect of context reconfiguration contrasted with subordinate rule reconfiguration in MFG and IFS. No significant clusters for the main effect of subordinate rule contrasted with stay (rule repeat) trials. **C)** Main effect of perceptual change (i.e., number of cue object features changing across trials; range 0-3) showing greater pattern shifts as more cue features changed across trials regardless of rule repeat/switch. **D-E**) Interactions between perceptual change and (**E**) subordinate rule reconfiguration or (**F**) rule repeat (Stay) trials showing distributed clusters with greater pattern shifts as more cue features changed. No significant clusters for the interaction between perceptual change and context reconfiguration. **F)** Perceptual change slopes for the 3 hierarchical repeat/switch conditions (Stay/IDS/EDS) in the MFG/IFS cluster (defined using the Schaefer 900 ROIs atlas; ROIs 780, 786, 789, & 793). Significantly positive slopes for Stay and IDS, but not significantly different from zero for EDS. **G)** Perceptual change slopes for each hierarchical conditions (Stay/IDS/EDS) in the caudal PFC cluster (Schaefer 900ROIs, 7 networks atlas; ROI 797), showing significantly positive slopes for Stay and IDS, but not for EDS. *** means p < .001, ** means p < .01, * means p < .05, n.s. means p >.05.

In line with these predictions, we identified two significant clusters in lateral frontal regions showing greater pattern shifts for EDS compared to IDS (inferior frontal sulcus (IFS) and caudal middle frontal gyrus (cMFG); Fig. 4B). Note, there were no significant clusters for the contrast between IDS and Stay trials. One possibility is that this null effect reflects overlap between this contrast and perceptual change, as well as our inclusion of the task switch control variable in the model. To address this, we conducted a supplementary analysis excluding perceptual change and task switch from the model (Fig. S1). This analysis revealed a significant caudal PFC cluster, located superior and posterior to the MFG cluster shown in Fig. 4B, that exhibited significantly greater pattern shift for IDS relative to Stay trials. Together, these results suggest that in our paradigm, pattern shift differences between IDS and Stay are primarily driven by perceptual change and task-switch related processes.

In contrast to the selective effect of context switches, perceptual change exerted widespread influences on multivoxel pattern shifts. A significant main effect of the number of changing cue features was observed across distributed regions (Fig. 4C), along with condition-specific simple effects of perceptual change within IDS trials (Fig. 4D) and Stay trials (Fig. 4E). We did not find any significant clusters for perceptual change within EDS trials. Within these regions, pattern shifts were greater when both task-relevant and task-irrelevant features changed compared to when only the task-relevant feature changed. These results are consistent with our hypothesis of increased susceptibility to perceptual interference during subordinate rule reconfiguration. These clusters were localized in a widely distributed fashion.

To ensure that the primary effects reported above were not driven by factors other than hierarchical switch type (Stay/IDS/EDS) and perceptual change, we examined several control variables included in the model. Consistent with the response design (right index/middle finger presses), response switches were associated with greater pattern shifts in left motor and visual cortices (Fig. S2A), reflecting this right-hand response design. Probe switches elicited greater pattern shifts in ventral occipito-temporal regions, including fusiform face area (FFA) and parahippocampal place area (PPA; Fig. S2B), consistent with the use of face and place stimuli. Previous target feature switches were associated with distributed effects spanning caudal PFC, visual cortex, medial temporal lobe, and basal ganglia (Fig. S2C). Post-error trials showed increased pattern shifts across medial prefrontal, medial temporal, and subcortical regions, including basal ganglia and mediodorsal thalamus (Fig. S3). No significant clusters were observed for the main effect of task switch, error trials, or the task × response interaction. We do not consider these control effects further as they do not bear directly on the hypotheses of interest.

To more precisely characterize the interaction between perceptual change and hierarchical task structure, we conducted two unbiased ROI analyses focusing on contrasting condition-specific effects of perceptual change within each hierarchical condition (Fig. 4F and G). This approach allowed us to directly test the predicted dissociation: perceptual change should have minimal impact during context switches (EDS) but modulate pattern shifts during subordinate rule switches (IDS) and rule repetitions (Stay). We defined two sets of ROIs based on independent whole-brain contrasts. First, to examine context-related effects, we selected ROIs within the lateral IFS-MFG ROIs overlapping with our searchlight cluster showing greater pattern shifts for EDS than IDS. Using the Schaefer 900-parcel (7-network) atlas, this cluster corresponded to four parcels (ROIs 780, 786, 789, and 793). Second, to show the specificity of this context switch effect in MFG, we selected an adjacent caudal MFG ROI that showed a main effect of perceptual change (ROI 797 in the same atlas).

Within the lateral IFS-MFG cluster, we extracted mean pattern shifts from these ROIs and applied the same pattern shift model as before, but in a mixed effects model with random intercepts for subjects. Similar to our z-scored RT results, we found positive slopes for the number of features changing across trials (perceptual change) for both rule repeat (Stay; *β* = 0.009, *SE* = 0.002, *CI* = [0.006, 0.013]), and subordinate rules switch (IDS; *β* = 0.012, *SE* = 0.002, *CI* = [0.008, 0.017]) and no reliable effect for context switch (EDS; *β* = -0.0002, *SE* = 0.002, *CI* = [-0.004, 0.004]). Follow-up contrasts confirmed that pattern shifts increased with greater perceptual change for Stay (Stay; zero versus one feature change; *mean difference* = - 0.009, *t(20965)* = -2.792, *p* = .032), and IDS (one versus two features change; *mean difference* = -0.012, *t(20957)* = -2.878, *p* = .024; Bonferroni corrected; Fig. 4F). In contrast, EDS trials showed no reliable differences across levels of perceptual change. Direct comparison of slopes further supported this dissociation, as the perceptual change slope was significantly more positive for IDS than EDS (*β* = 0.013, *t(20953)* = 2.394, *p* = .044). A post-hoc contrast between context switch and subordinate rule switch trials further confirmed that context switch trials show significantly larger pattern shifts than subordinate rule switch trials (*β* = 0.016, *t(20952)* = 2.829, *p* = .014).

We next examined the caudal MFG ROI defined from the main effect of perceptual change (Fig 4G). Using the same modeling approach, we again observed positive slopes for perceptual change in Stay (*β* = 0.010, *SE* = 0.004, *CI* = [0.004, 0.017]) and IDS trials (*β* = 0.015, *SE* = 0.005, *CI* = [0.006, 0.023]), with no reliable effect for EDS trials (*β* = 0.002, *SE* = 0.004, *CI* = [-0.005, 0.010]). Follow-up contrasts replicated this pattern of slope effects: pattern shifts increased with greater perceptual change for Stay (zero versus one feature; *mean difference* = - 0.010, *z* = -2.982, *p* = .017) and IDS trials (one versus two features change; *mean difference* = - 0.015, *z* = -3.256, *p* = .007), but not for EDS trials. However, slopes did not differ between hierarchical conditions (*β* = 0.013, *t(20938)* = 0.756, *p* = .730). Moreover, we confirmed a lack of a context switch effect (*β* = 0.009, *t(20956)* = 1.649, *p* = .297) for this more caudal PFC ROI.

These multivoxel pattern shifts results support our hypotheses on cognitive flexibility. Perceptual change modulates neural representations differently depending on the level of hierarchical representations. During subordinate rule switches (IDS), pattern shifts increase with the number of changing cue features, indicating greater sensitivity to task-irrelevant perceptual information. In contrast, context switches (EDS) elicit larger overall multivoxel pattern shifts that are invariant to perceptual change, consistent with more abstract representations that are better insulated from feedforward interference.

### 3.3. Conjunctive representations supporting cognitive flexibility are context dependent

Prior work suggests cognitive flexibility, i.e., fast and efficient switching between rules/tasks, is supported by the creation of conjunctive representations (Kikumoto and Mayr, 2020). These conjunctive representations consist of task rules, stimuli, and responses. A common measure of conjunctions has been partial switch costs—greater switch costs when only a subset of conjunction components change across trials, as compared to trials where all components switch or repeat (Grzyb & Hubner, 2013; Kleinsorge & Heuer, 1999). When all components of a conjunction repeat, then the previous trial’s conjunction can be re-instantiated. However, when some components repeat while others switch, the previous conjunction needs to first be unbound so that a new conjunction can be created with the components that repeat across trials. Furthermore, the conjunction components that repeat across trials can cause increased interference when sensory-motor plans differ between these two trials. Thus, this unbinding and rebinding in addition to increased interference that needs to be resolved cause the observed “cost’. When all components of a conjunction switch, the unbinding process can be skipped and the lack of shared components with the previous trial means no additional interference needs to be overcome. However, a new conjunction still needs to be constructed, which is why response times are still slower overall than trials where the previous conjunction can be re-instated and utilized.

Previous work on conjunctive representations has largely examined tasks with a single level rule structure (Kikumoto and Mayr, 2020; Rangel et al., 2023). As such, it remains an open question as to whether conjunctions consist of only the task sets organizing sensory-motor plans, or if they include the higher-order context-level information that our paradigm was designed to probe. Recent evidence suggests that conjunctive representations may be organized into context-specific subspaces, enabling more efficient readout of task-relevant information (Kikumoto et al., 2024). Given that in our task, context indicates if color or shape is the currently relevant rule, we predicted that conjunction-related partial switch costs would be observed within each context, but not when switching across contexts. Under this account, context serves to index distinct conjunctive subspaces, such that switching context both circumvents the need to unbind the previous conjunction and reduces interference from the previous conjunction.

To measure this, we first ran a modified version of the above RT mixed effects model where the hierarchical rule switch (Stay/IDS/EDS) was replaced with a variable indicating if context switched or repeated (2 levels). This allowed us to run a three-way interaction between context, task, and response, while controlling for all the same additional variables as before. We found a significant three-way interaction (*β* = 0.128, *t(20930)* = 2.142, *p* = .032). Post-hoc comparisons revealed that the two-way interaction between task and response, which reflects partial switch costs, was only significant when context repeated (*β* = -0.124, *t(20916)* = -2.300, *p* = .022; Fig. 5A), but not when context switched (*β* = 0.004, *t(20926)* = 0.065, *p* = .949; Fig. 5B). To replicate this result, we ran a follow-up behavioral experiment with the same experimental paradigm. As expected, we found a significant three-way interaction (*β* = 0.251, *t(21690)* = 4.173, *p* < .001). Post-hoc comparisons once again revealed that the two-way interaction between task and response, which reflects partial switch costs, was only significant when context repeated (*β* = -0.247, *t(21704)* = -4.891, *p* < .001; Fig. 5C), but not when context switched (*β* = 0.004, *t(21687)* = 0.056, *p* = .955; Fig. 5D).

**Figure 5.**
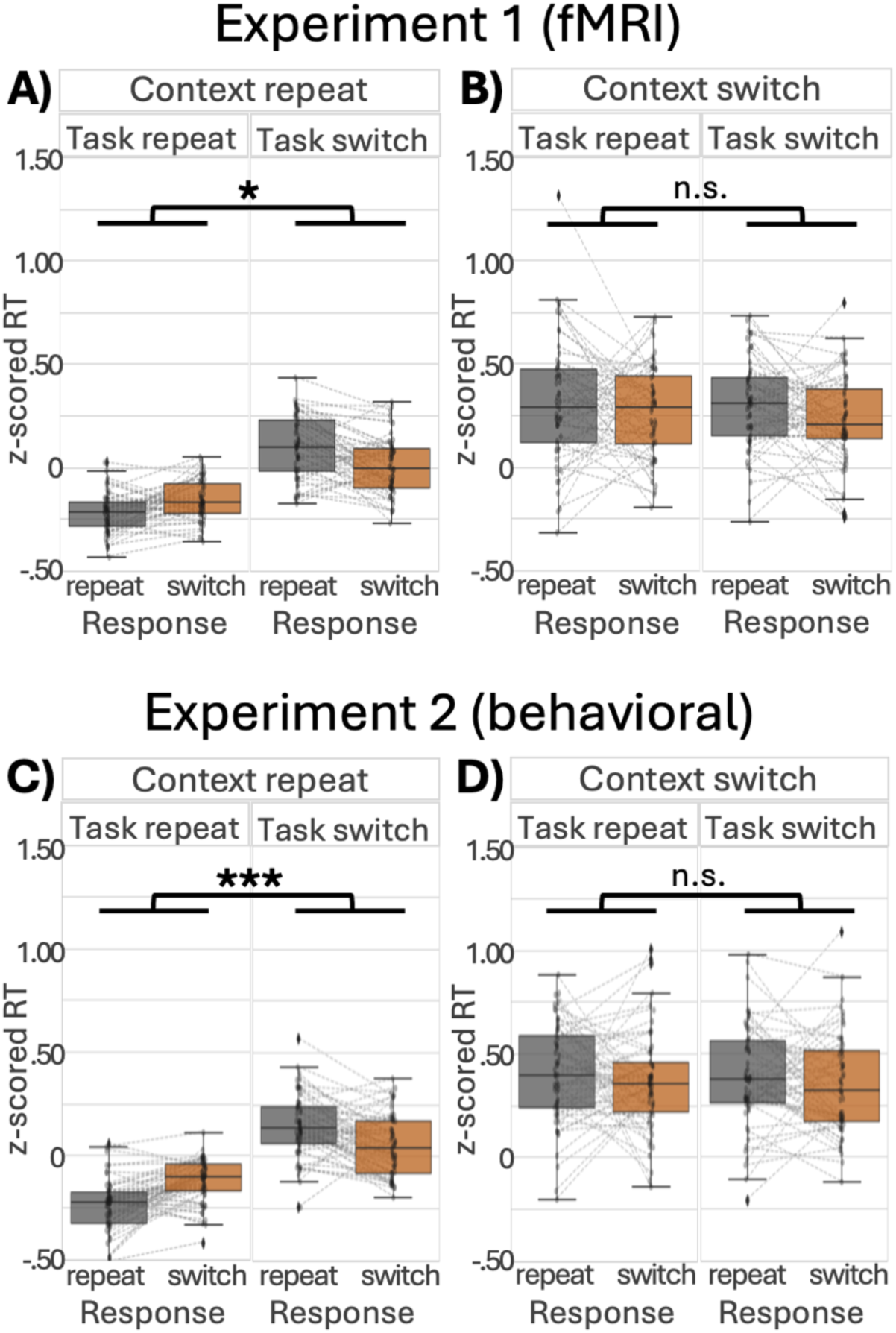
Partial Switch Costs and Context Reconfiguration. **A&C**) Bar plots showing expected partial switch cost effects when context repeats. **B&D**) Bar plots showing lack of partial switch costs when context switches. Significance based on post-hoc comparisons between estimated marginal means for 3way interaction between context, task, and response repeat/switch. *** means p < .001, ** means p < .01, * means p < .05, n.s. means p >.05.

The existence of a significant two-way interaction between task and response within context matches prior findings of partial switch costs (Kikumoto and Mayr, 2020; Rangel et al., 2023). In contrast, the lack of a significant two-way interaction when switching contexts suggests that partial switch costs are not observed during context reconfiguration. These results lend further support to the idea that cognitive flexibility is supported by creating context-specific conjunctive representations, which enable faster subordinate rule switching within context and decreased interference when switching across context.

To address the neural underpinnings of partial switch costs indicative of potential context-specific conjunctive representations, we conducted a parallel analysis with a very similar regression model to predict multivoxel pattern shifts. Key differences were that cue color, shape, and texture were removed (similar to the regression predicting pattern shifts reported in the previous results section) and regressors indicating the z-scored RT and its derivative were included. However, we found no significant clusters associated with a three-way interaction between context, task, and response repeat/switch. This suggests that, although the observed partial switch cost effect on context repeat trials, but not context switch trials, is robust for response times, its neural underpinnings likely arise from a combination of distributed neural representations rather than from anatomically focal, restricted sources.

## 4. Discussion

In this study, we investigated the behavioral (RT) and neural correlates of switch costs associated with hierarchical cognitive control. We were primarily interested in how the reconfiguration of distinct cognitive representations would interact with (1) the underlying hierarchical task representation structure and (2) perceptual systems/processes necessary for goal-directed behavior. To address these questions, we combined a hierarchical control task with trial-by-trial behavioral switch cost analyses and a multivariate distance measure on voxel patterns (“pattern shifts”), allowing us to characterize both performance costs and representational reconfiguration during hierarchical cognitive control. In the sections that follow, we interpret these findings in the context of existing theories of cognitive control and representational flexibility.

The behavioral costs associated with task switching is well-established (Allport, Styles, & Hsieh, 1994; Monsell, 2003; Rogers & Monsell, 1995; Wylie & Allport, 2000). However, less is known about their neural basis, particularly how they relate to representational structure. A previous study found that greater dissimilarity in multivoxel activity patterns on task switch trials predicted greater RT switch costs (Qiao et al., 2017). Nevertheless, given the multifaceted nature of task representations (Botvinick, 2008; Schumacher & Hazeltine, 2016), several critical questions remain unresolved: whether representational reconfiguration operates similarly across different hierarchical levels, how abstract contextual representations interact with perceptual inputs, and whether perceptual processes shape reconfiguration in ways that support cognitive flexibility. To address these questions, we extended multivoxel pattern shift analyses to a hierarchical task structure that dissociates contextual updating from subordinate rule reconfiguration, allowing us to test how different representational levels vary in their susceptibility to perceptual interference.

A complementary line of work suggests that stimulus-response associations are bound into an event file (Hommel, 1998, 2004), or task-rule conjunctions (Kikumoto & Mayr, 2020). Reusing these integrated sensory-motor representations facilitates performance; if updated, performance costs increase, particularly under partial overlap between trial components (Grzyb & Hubner, 2013; Kleinsorge & Heuer, 1999). Despite this, relatively little work has examined how conjunctive representations operate within hierarchical task structures. Kikumoto and colleagues (2024) found evidence for context-specific conjunctions, however, context served as a retro-cue for conjunction selection. Thus, it remains unclear whether conjunctions incorporate context when context and rule information are presented concurrently. To address this question, we examined how hierarchical context, feedforward perceptual inputs, and conjunctive representations might interact to shape behavioral and neural switch costs.

In our task, rule/task set selection was shaped by both top-down contextual modulation and bottom-up perceptual inputs. Critically, we found that context reconfiguration changed the balance between top-down and bottom-up influences. Subordinate rule reconfiguration was more susceptible to feedforward perceptual interference than context reconfiguration, suggesting bottom-up influence on task-set selection within a stable context. Task-irrelevant feature changes slowed RTs and increased pattern shifts across distributed frontal, premotor, and occipitotemporal regions. In contrast, context reconfiguration showed little modulation by task-irrelevant feature changes in MFG, with similarly large RT costs and pattern shifts regardless of perceptual overlap. Given work suggesting MFG encodes high dimensional/abstract task rule structures (Badre et al., 2021; Badre & Nee, 2018), reduced susceptibility to interference is critical for successful goal-directed behavior.

Importantly, differential susceptibility to perceptual interference should not be interpreted as independence between hierarchical levels. Classical hierarchical accounts primarily characterize top-down relationships, whereby higher-order contextual representations influence subordinate rule selection (Badre et al., 2009; Koechlin, Ody, & Kouneiher, 2003). Our study, however, examined the influence of bottom-up, task-irrelevant perceptual inputs on reconfiguration processes. While the hierarchical framework describes how higher-level representations guide lower-level processing, they do not predict equal susceptibility to feedforward interference at all levels. Our findings are novel in suggesting that contextual representations can guide subordinate rule selection while remaining relatively insulated from irrelevant perceptual inputs, potentially reflecting a mechanism that preserves stable goal representations during hierarchical reconfiguration.

Contextual representations might reduce feedforward interference by organizing lower-level perceptual/motor representations into distinct representational subspaces. Kikumoto and colleagues (2024) suggest the control system encodes conjunctive representations (integrated representations of task rules, stimuli, and responses) in distinct subspaces. Context-specific conjunctions would allow for a smaller representational distance within context, meaning greater flexibility, while also shielding current goal-relevant conjunctions from interference across contexts. Greater flexibility within context likely contributes to observed partial switch costs when context repeats. In contrast, shifting contextual subspaces requires traversing larger representational distances, potentially explaining slower RTs and the lack of partial switch costs when context changes.

While our framework suggests context representations reduce feedforward interference, an alternative account is that interference from task-irrelevant features still occurs during context reconfiguration but unfolds in parallel with reconfiguration processes. If context updating is the slower process, it could obscure behavioral cost arising from perceptual interference. This account would predict a modulation of multivoxel pattern shifts by task-irrelevant features across all hierarchical conditions. We found that, although both Stay and IDS trials showed increased pattern shifts with task-irrelevant feature changes across distributed regions, no such effect was observed for EDS trials in any region. Therefore, these results do not support the parallel processing account and instead support our argument that contextual representations are relatively insulated from feedforward perceptual interference.

The present findings raise a central question: What mechanism drives the observed pattern shifts and supports the updating of hierarchical task representations? Converging evidence from our prior work and others (Chen et al., 2024; Mukherjee et al., 2021; Rikhye, Gilra, & Halassa, 2018) points to the mediodorsal thalamus as a critical subcortical region in this process. Previously, we found that the mediodorsal thalamus is more strongly engaged during context updating (Chen et al., 2024). Building on this finding, the current results suggest that thalamocortical interactions may regulate the extent to which feedforward perceptual inputs influence the updating of context/context-specific conjunctive representations. In this way, thalamocortical interactions may enable flexible reconfiguration while limiting interference from task-irrelevant inputs.

One possibility is that the mediodorsal thalamus mediates a context-specific updating signal that implements a form of input gating (Rac-Lubashevsky & Frank, 2021). During context reconfiguration, thalamocortical interactions mediated by mediodorsal thalamus could drive targeted updating of cortical context representations while constraining the influence of competing feedforward inputs. Although we did not observe significant corrected clusters in mediodorsal thalamus during EDS trials in the present study, this account is supported by our prior findings, where the mediodorsal thalamus showed increased evoked responses and functional connectivity during context switches (Chen et al., 2024). Under this view, context updating may involve selectively admitting context-relevant signals while restricting task-irrelevant sensory inputs from propagating into higher-order representations. Such gating could be implemented via known circuits, including basal ganglia–mediated control of thalamic relay activity (e.g., input/output gating; (Chatham, Frank, & Badre, 2014; O’Reilly & Frank, 2006) or inhibiting feedforward inputs through the thalamic reticular nucleus (Halassa & Acsady, 2016; Zikopoulos & Barbas, 2006). However, the present fMRI data do not allow for distinguishing between these possibilities. During IDS and Stay trials, when context is stable, this gating may be relaxed or differently configured, allowing feedforward inputs to more directly influence rule and conjunctive representations. This would permit multiple feature inputs to contribute to the formation and updating of conjunctive representations, consistent with the distributed cortical effects of perceptual change observed here, as well as the involvement of posterior thalamic regions (Fig. 4C–D).

Our measure of multivoxel pattern shifts has limitations that need to be addressed. First, while greater dissimilarity in multivoxel activity patterns implies less similar population responses, it can also be driven by multiple coding geometries (e.g., orthogonal subspaces versus shifts within a shared space). Our measure detects the net effect of representational changes influencing voxel patterns but does not uniquely specify the underlying neuronal population geometry beyond the resolution of fMRI. Distinct task variables may coexist within voxels and may be implemented as partially separable subspaces within the same population within the same voxel (Weber et al., 2023). Moreover, according to Walther (2016), pattern shifts estimated with correlation distance can be obscured when conditions share high overall magnitude shifts. Future work could evaluate the extent to which the observed effects generalize across alternative distance metrics, such as Mahalanobis distance.

In conclusion, we report three major findings regarding switch costs and hierarchical cognitive control. First, trial-to-trial reconfiguration of neural activity patterns drive observable switch costs, with greater pattern reconfiguration predicting larger switch costs. Second, the reconfiguration of distinct hierarchical levels is differentially impacted by feedforward perceptual information, as is evident through both RT and pattern shift measures. Specifically, subordinate rule representations associated with more concrete sensory-motor plans prioritized greater flexibility at the cost of greater feedforward perceptual interference. To counter this increased interference, contextual representations, which are more abstracted away from immediate sensory-motor plans, shielded information from feedforward perceptual interference at the cost of reduced flexibility. This likely reflects a balance of cognitive flexibility that supports faster adaptation of concrete action plans while shielding internally maintained contextual representations from interference. Finally, RT switch cost results suggest that this shielding is achieved by partitioning contexts into distinct representational subspaces and forming context-specific conjunctions within these subspaces.

## Acknowledgments

Author contributions: SCL wrote the first draft of the paper, SCL and KH edited the paper, KH and SCL designed research, SCL and XC performed research, SCL and KH analyzed data.

## Authors declare that they have no competing interest

All data and code will be made available upon publication.

Research reported here was supported by a National Institutes of Mental Health grant R01MH122613 (KH) and the Iowa Neuroscience Institute. This work was conducted on an MRI instrument funded by 1S10OD025025-01

