## supplement for "Hierarchical Reconfiguration of Neurocognitive Task Set Representations Mediates Cognitive Flexibility"

Stephanie C. Leach* *et al.*

**This PDF file includes:**

Figures S1-S3

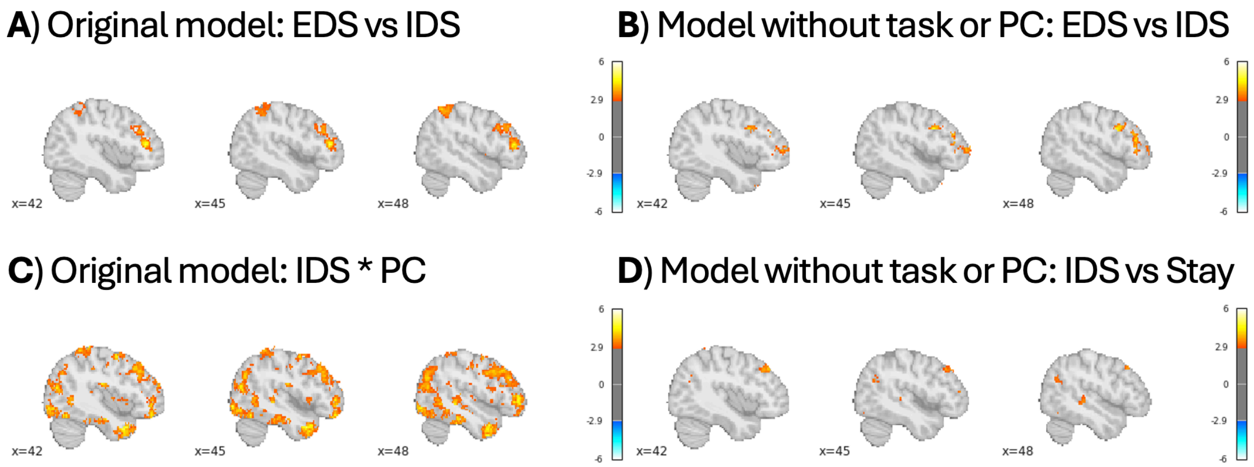


**Figure S1**. **Supplementary Analysis of Pattern Shifts without Task and Perceptual Change Control Variables**. **A**) Significant clusters associated with a larger pattern shift on context switch (EDS) versus subordinate rule switch (IDS) trials based on the original model reported in the main manuscript. **B**) Significant clusters associated with a larger pattern shift on context switch (EDS) versus subordinate rule switch (IDS) trials based on a supplementary model excluding task and perceptual change (PC) from the list of predictor variables. **C**) Significant clusters associated with a larger pattern shift when more cue object features change on subordinate rule switch (IDS) trials based on the original model reported in the main manuscript. **D**) Significant clusters with a larger pattern shift on subordinate rule switch (IDS) versus rule repeat (Stay) trials based on a supplementary model excluding task and perceptual change (PC) from the list of predictor variables.

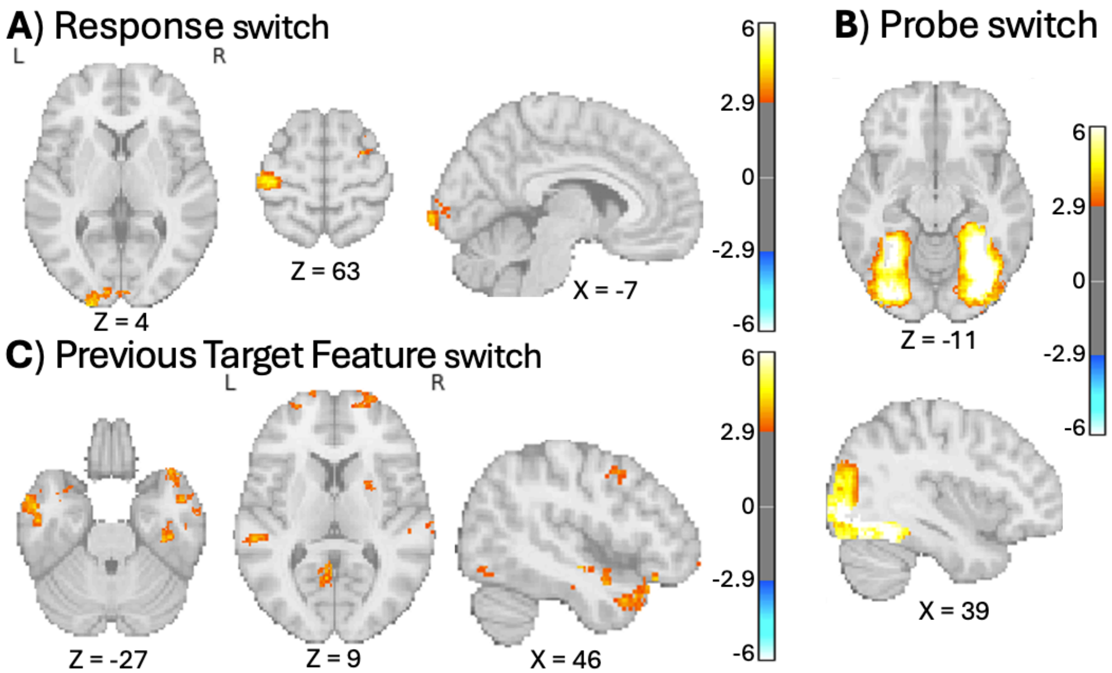


**Figure S2**. **Task, Response, and Previous Target Feature Switch versus Repeat.** A) main effect of response reconfiguration contrasted with response repeat—significant clusters in left primary motor cortex and visual cortex. B) main effect of probe image switch contrasted with probe image repeat—significant clusters in ventral visual stream (including FFA/PPA). C) main effect of previous target feature switch contrasted with previous target feature repeat—notably, significant clusters in caudal PFC, primary visual cortex, right parahippocampus to inferior and middle temporal gyrus, and the right putamen.

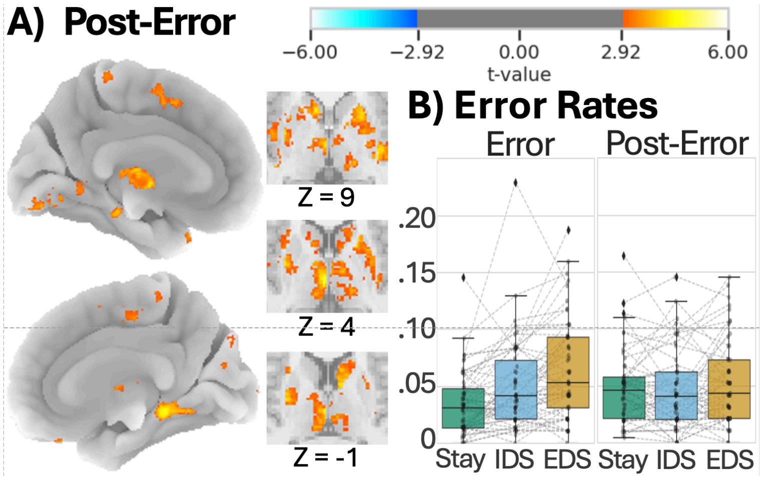


**Figure S3**. **Errors and Post-error Correction**. A) Significant clusters with a larger pattern shift on post-error trials B) Error and Post-error trial rates by hierarchical rule condition (Stay/IDS/EDS).
